# Functional traits of ectomycorrhizal trees influence their effects on surrounding soil organic matter properties

**DOI:** 10.1101/2024.12.13.628221

**Authors:** Joseph D. Edwards, James W. Dalling, Jennifer M. Fraterrigo, William C. Eddy, Wendy H Yang

## Abstract

Ectomycorrhizal (EM) effects on forest ecosystem carbon (C) and nitrogen (N) cycling are highly variable, which may be due to underappreciated functional differences among EM-associating trees. We hypothesize that differences in functional traits among EM tree genera will correspond to differences in soil organic matter (SOM) dynamics.

We explored how differences among three genera of angiosperm EM trees (*Quercus, Carya,* and *Tilia*) in functional traits associated with leaf litter quality, resource use and allocation patterns, and microbiome assembly related to overall soil biogeochemical properties.

In support of our hypothesis, we found consistent differences among EM tree genera in function traits. *Quercus* trees had lower litter quality, lower δ^13^C in SOM, higher δ^15^N in leaf tissues, greater oxidative extracellular enzyme activities, and higher EM fungal diversity than *Tilia* trees, while *Carya* trees were often intermediary. These functional traits corresponded to overall SOM C and N dynamics and soil fungal and bacterial community composition.

Our findings suggest that trait variation among EM-associating tree species should be an important consideration in assessing plant-soil relationships such that EM trees cannot be categorized as a unified functional guild.

## Introduction

Symbiosis between plants and microbes are critical for surrounding ecosystem function, helping mediate organismal fitness (Benning and Moeller, 2021), biogeochemical cycling (Rousk and Bengtson, 2014), and resilience to global change (van der Putten et al., 2016). We often classify various organisms into broader functional groups based on shared traits or phylogenies to help explain their role in the environment. However, these functional groups are often too broad, ignoring potentially important diversity among organisms with other shared traits (Taylor et al., 2020, Love et al., 2023). For example, mycorrhizal associations between trees and fungi are increasingly used as a tool to predict forest nutrient cycling and soil organic matter (SOM) turnover (Phillips et al., 2013) as well as potential responses to global change (Terrer et al., 2016, Sulman et al., 2019). In these mycorrhizal frameworks, forests are classified as either ectomycorrhizal (EM) or arbuscular mycorrhizal associating based on the species identity of dominant trees, but these two groups may not be wholly comparable. Ectomycorrhizal associating trees are a phylogenetically diverse group (Averill et al., 2019, Tedersoo et al., 2010) with significant functional diversity in important ecosystem traits like SOM accumulation (Hicks Pries et al., 2023), decomposition (Fernandez et al., 2019), and partner specificity (Anthony et al., 2022). These functional differences may be responsible for the considerable variation in EM-associated effects on ecosystem, particularly carbon (C) and nitrogen (N) cycling (Fernandez and Kennedy, 2016, Lin et al., 2017). Yet, we still know relatively little about the mechanisms and functional traits that drive divergent ecosystem responses to different EM-associating trees.

Plant-soil feedbacks that contribute to EM-associated ecosystem effects are driven by both plant and microbial mechanisms (Herrera Paredes and Lebeis, 2016, Van der Putten et al., 2013). In the context of nutrient cycling and SOM dynamics, the most important of these mechanisms likely relate to how resources are being acquired and allocated in plants and soils via (1) leaf litter quality, (2) resource acquisition and allocation patterns, and (3) microbiome assembly and function (Figure 1).

*1. Leaf litter quality*: From a plant perspective, EM trees often have nutrient-conservative traits with high foliar C : nutrient stoichiometries (Averill et al., 2019); however, angiosperm EM trees can exhibit up to two-fold differences in litter quality (Craig et al., 2018). These differences in leaf litter quality can influence the strength of EM effects on SOM dynamics (Fernandez et al., 2019, Seyfried et al., 2021). However, litter quality does not fully explain differences in decomposition rates among EM-dominated stands (Midgley et al., 2015), suggesting an important role for microbial factors in mediating this relationship.
*2. Resource acquisition and allocation patterns*: Ectomycorrhizal trees are generally assumed to invest more in their fungal partners than non-EM trees (Terrer et al., 2016), however, belowground C allocation can vary greatly among forest stands with differing EM tree community assemblages (Keller et al., 2021). As EM fungi can adjust how much N they allocate to hosts based on C investments (Bogar et al., 2022, Horning et al., 2023), these mutualism resource allocation and acquisition strategies may lead to downstream effects on soil communities and biogeochemistry. Greater belowground C allocation from trees can increase EM fungal biomass in soils (Hobbie, 2006), promoting the accumulation of soil C as these organisms turnover and necromass is converted into more stable SOM (Hu et al., 2023), such as mineral-associated organic matter (MAOM; Lavallee et al., 2019, Klink et al., 2022). Greater N acquisition by EM fungi could promote increased competition with saprotrophic fungi, decreasing decomposition in favor of soil C accumulation (Bödeker et al., 2016).
*3. Microbiome assembly and function*: Differences among EM trees in their mutualist and free-living microbiome assembly based on their relatively high partner specificity (Lofgren et al., 2018) could also drive variation in their plant-soil feedbacks. Ectomycorrhizal fungi often liberate nutrients from SOM via oxidative enzymes (Shah et al., 2016) with these extracellular enzyme activities (EEAs) differing among EM fungal lineages (Pellitier and Zak, 2018). Further, unique competition and cooperation dynamics may emerge between EM fungi and free-living microbiomes (Edwards et al., 2024, Eagar et al., 2024) based on EM fungal identity and function (Zak et al., 2019). To assess the importance of tree identity in mediating EM-associated plant-soil feedbacks, we must understand the relationship between their functional traits (*i.e.,* litter quality, resource use patterns, and microbiome assembly) and effects on ecosystem function.

**Figure 1.**
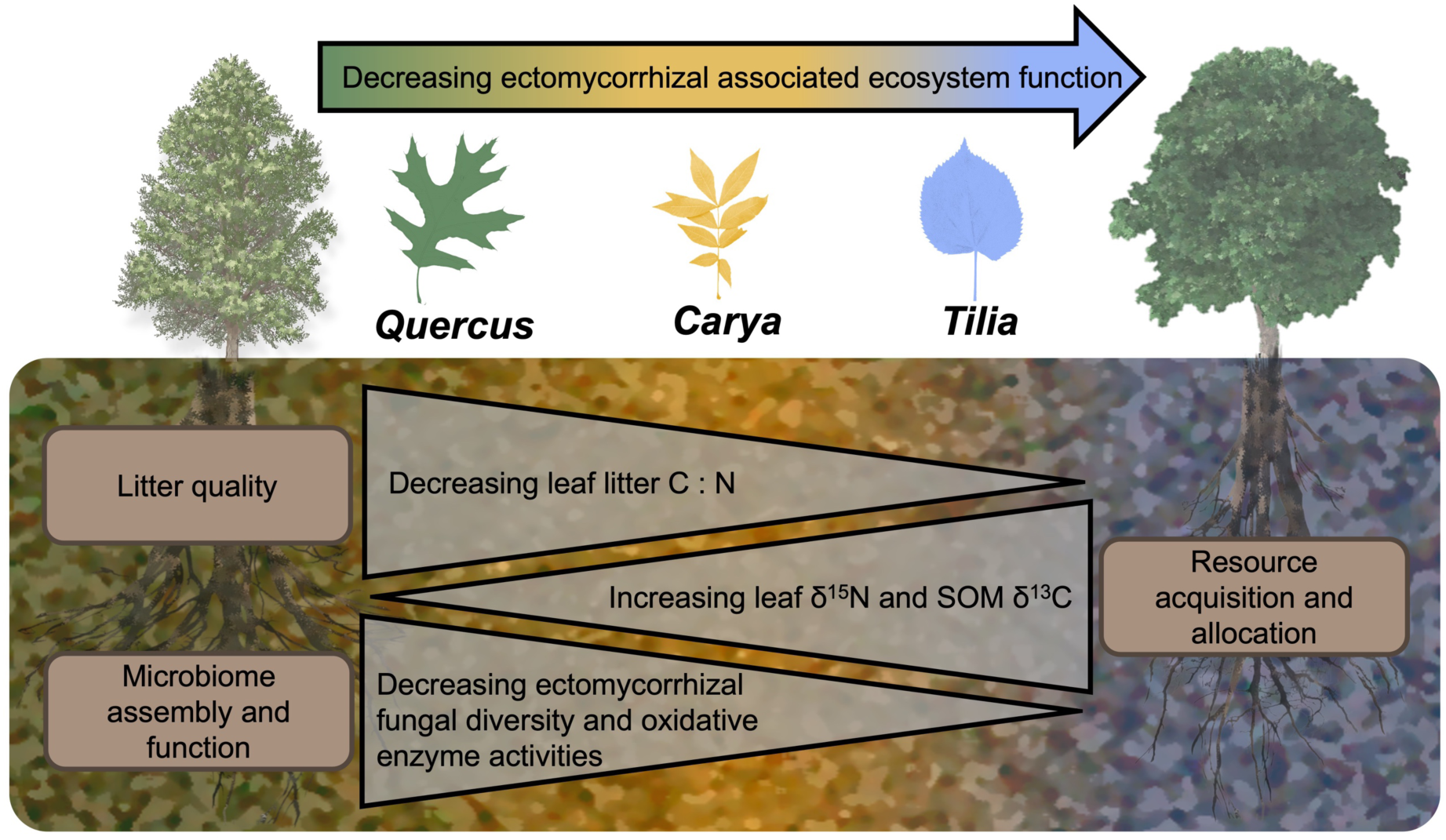
Conceptual diagram showing hypothesized relationship between ectomycorrhizal- associated ecosystem function and EM functional traits driven by decreasing litter quality, changes to resource acquisition and allocation, and shifts in microbiome assembly and function.

We hypothesize that variation in observed in EM effects on SOM and surrounding ecosystems is linked to the identity of EM-association trees. Therefore, differences in functional traits among EM tree genera will correspond to differences in soil biogeochemical properties (Figure 1). We assessed these relationships for three types of functional traits thought to be important for EM symbiosis: 1) leaf litter quality, assessed by C : N stoichiometry; 2) resource use patterns, assessed by natural abundance C and N stable isotopic composition of leaf litter and SOM; and 3) EM fungal and free-living microbial community assembly, assessed by the diversity and composition EM fungal, general fungal, and bacterial communities. Elucidating these patterns and drivers of EM-associated plant-soil feedbacks and SOM properties will improve our ability to leverage forest stand mycorrhizal types and community composition to understand and predict their effects on surrounding ecosystems.

## Methods

To test the hypotheses that different effects of EM trees on surrounding ecosystems are mediated by their functional traits, we sampled soils and leaf litter from near three different EM tree genera with known overall functional differences (*Carya, Quercus,* and *Tilia*) in an EM dense forest patch in central Illinois, USA. We conducted our study in Trelease Woods, a 24 ha remnant old-growth mixed mesophytic forest near Urbana, IL, USA (44.13°N, -88.14°W). Mean annual temperate at this site is 10.5 °C and mean annual precipitation is 1010 mm. Census data from 2018 were used to identify a 200m x 175m area dominated by EM-associating trees of the genera *Carya* (*C. laciniosa, C. ovata*), *Quercus* (*Q. macrocarpa, Q. rubra*), and *Tilia* (*T. americana*; Figure S1). These tree genera differ in traits that may be related to their mycorrhizal function: litter production and chemistry (Peterson and Cummins, 1974, CÔtÉ and Fyles, 1994), root morphology (Beals and Cope, 1964, Brundrett et al., 1990), and foliar and reproductive phenology (Lechowicz, 1984). There is significant phylogenetic conservation for both collaborative and conservative traits among tree species (Bergmann et al., 2020); thus, we use genus as our central focal identification for EM trees in this study. We identified the 25 largest individual trees of each genus within the forest stand for sampling (Figure S1). Samples were collected from within 5 meters of each focal tree in a randomly chosen cardinal direction.

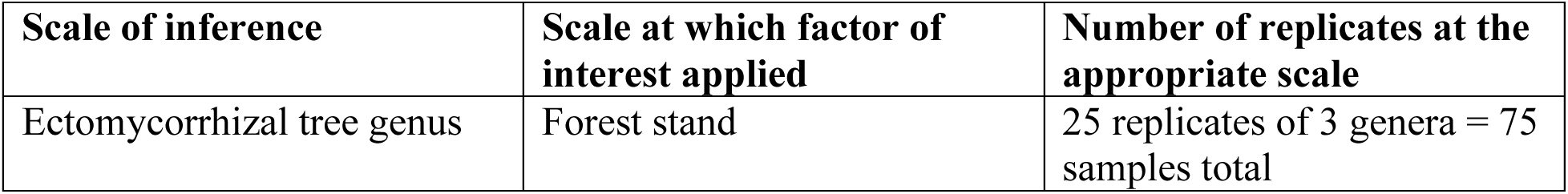

Soil samples were collected in late June of 2020 using a 5 cm diameter soil corer. Timing of soil sampling was based on previous estimates of seasonal maximum EM fungal activity/ relative abundance at this site (Edwards et al., 2022a, Edwards et al., 2022b). Soil samples were collected intact in a plastic sleeve to preserve root integrity. Samples were immediately capped on both ends and put on ice for transport to the University of Illinois at Urbana-Champaign later that day. Following collection, all samples were hand-picked to remove rocks, coarse litter/roots, and visible fine roots (< 2mm diameter), then sieved through a 2 mm mesh. Soils were then subsampled for the analyses for soil pH, moisture, inorganic N, dissolved organic C (DOC), total dissolved N (TDN), microbial biomass C (MBC), and microbial biomass N (MBN) and immediately processed. A separate subsample of soil was frozen at -20° C for later analysis of soil DNA, EEAs, SOM size fractions, and C and N elemental concentrations and isotopic composition. All samples were processed and frozen (if necessary) within 12 hours of collection. We also collected fresh litter from each focal tree (25 samples per species) using 50 cm by 30 cm litter traps in October – November 2021 because major defoliation from a hailstorm in August 2020 precluded litter collection in the same growing season as soil sampling. Leaf litter was collected daily from litter traps until sufficient to perform chemical analyses.

### Assessment of functional traits

We assess how litter quality, resource allocation and acquisition patterns, and microbiome assembly influenced genus-specific effects on EM trees on surrounding ecosystems.

*1. Litter quality:* Litter quality was measured using the C : N stoichiometry of leaf litter. Leaf litter quality is highest (low C:N, high pH, low lignin) in *Tilia* species (Page and Mitchell, 2008) and lowest in *Quercus* species (Scott and Binkley, 1997, Wilson et al., 2022), while *Carya* leaf litter quality is intermediate yet highly variable among species (CÔtÉ and Fyles, 1994, Vesterdal et al., 2008).
*2. Resource allocation and acquisition:* Resource acquisition and allocation patterns were assessed using the natural abundance stable isotopic composition of C and N in leaf litter and SOM. Stable isotopes are a powerful integrator of plant resource use as they reflect the net outcome of many processes (Dawson et al., 2002). The direct transfer of C and N between EM plants and fungi leads to a distinct isotopic signature in the biomass of these organisms relative to those that acquire resources from other sources (Taylor et al., 2003). Ectomycorrhizal fungal N transfer to plants discriminates against the heavier ^15^N isotopes, resulting in foliar ^15^N depletion (lower δ^15^N) in plants that rely more strongly on EM mutualists for N acquisition (Hobbie and Högberg, 2012). In contrast, EM fungi are more depleted in ^13^C (lower δ^13^C) than free-living saprotrophs because differences in the C isotopic signature of their different C sources, rhizodeposits and SOM, respectively (Hou et al., 2012, Hobbie et al., 2001). Since fungal necromass is a major contributor to MAOM in temperate forest soils (Klink et al., 2022), a lower C isotopic value for MAOM can reflect greater relative contribution of EM to MAOM formation than saprotrophic fungi. However, variation in N source pools or in C transfer from trees to EM fungi can confound these interpretations. We accounted for this by evaluating the difference in C and N isotopic composition between leaf litter and MAOM (*i.e.* δ^13^CMAOM-litter and δ^15^NMAOM-litter). The isotopic signature of N source pools, which can vary with soil depth and N form, imprints on the isotopic values for both the leaf litter and mycorrhizal fungal biomass such that greater mycorrhizal transfer of N would lead to a greater difference in δ^15^N between these two pools (Seyfried et al. 2021a). As the contribution of N acquired by mycorrhizal fungi to leaf litter increases, this signal of resource use and acquisition becomes stronger in δ^15^NMAOM- litter by causing this value to increase. In contrast, lower δ^15^NMAOM-litter could reflect either low rates of resource exchange between mycorrhizal fungi and their tree partners or low contributions of mycorrhizal necromass to MAOM formation. The isotopic signature of C fixed by trees is similarly imprinted in leaf litter and in the photosynthate transferred belowground to roots, rhizodeposits, and mycorrhizal fungi. In this way, a smaller δ^13^CMAOM-litter could reflect greater resemblance between these two pools as trees increase belowground C allocation via root growth, rhizodeposition, or mycorrhizal C transfer. Therefore, differences among tree genera in their δ^13^CMAOM-litter and δ^15^NMAOM-litter can reflect biological processes related to the acquisition and allocation of these resources. Natural abundance C and N isotopic composition of leaf litter and SOM can also be influenced by other environmental mechanisms (Craine et al., 2009). Many of the environmental gradients promoting differences in isotopic composition (e.g., precipitation, parent material, N deposition) occur at large spatial scales, and thus are minimized by our local-scale approach. We also empirically accounted for other potential influences on C and N isotopic composition of leaf litter and SOM in two ways. First, we incorporated spatial autocorrelation into statistical tests to account for spatial heterogeneity in isotopic signatures of C and N sources (see statistical analysis section). Second, we assessed variables related to soil fertility that could influence leaf litter δ^15^N by altering plant N uptake patterns, such as soil acid-base chemistry and inorganic N concentrations (Lin et al., 2021).
*3. Microbiome assembly and function:* Differences in microbiome assembly among EM tree genera were assessed via EM fungal and free-living fungal and bacterial communities alpha and beta diversity. Microbiome function was assessed by major oxidative and hydrolytic EEAs for liberating nutrients and metabolizing C from SOM.

### Soil and leaf litter properties

To assess differences in leaf and soil traits among EM tree genera, we measured chemical and physical properties of leaf litter, bulk soils, and SOM pools. To partition mineral-associated and particulate organic matter contributions to overall SOM, we used size fractionation following Bradford et al. (2008). Briefly, soils were shaken in 5% Sodium hexametaphosphate for ∼18 hours, then sieved through 53 µm mesh. Sample that passed through the mesh was considered MAOM, and sample caught by the mesh was considered particulate organic matter (POM). Both size fractions were collected and dried at 65 °C. Bulk soil, SOM fractions, and litter samples were dried and pulverized via bead-beating before being analyzed for C and N elemental concentrations and isotopic composition using an Vario Micro Cube elemental analyzer (Elementar, Hanau, Germany) interfaced with an IsoPrime 100 isotope ratio mass spectrometer (Isoprime Ltd., Cheadle Hulme, UK). Analytical precision was measured based on the standard error of calibrated QC standards (n = 14) run across all batches. The analytical precision of δ^13^C was 0.044 ‰ and the analytical precision of δ^15^N was 0.086 ‰.

Gravimetric soil moisture was measured by oven-drying another subsample of each soil at 105 °C for 24 hours. Soil pH was determined using a 1:1 mass ratio of soil:DI H2O. To measure soil inorganic N concentrations, soils were extracted in 2M KCl, and soil NO3^-^ and NH4^+^ concentrations were determined by analyzing the extracts using colorimetric methods on a SmartChem 200 discrete analyzer (KPM analytics, Westborough, MA). To measure dissolved organic C (DOC), total dissolved N (TDN), microbial biomass C (MBC), and microbial biomass N (MBN), soils were extracted in 0.05M K2SO4 with and without a direct chloroform addition (Brookes et al., 1985). Non-purgeable organic C and total N concentrations in the extracts were determined on a Shimadzu TOC-V CSH (Shimadzu Corp., Kyoto, Japan). Microbial extracted organic C and N from chloroform-fumigated samples was converted to MBC and MBN using an assumed extraction efficiency of 0.17 (Gregorich et al., 1990), based on the relatively low concentration of K2SO4. Base cation concentrations and cation exchange capacity (CEC) were measured by extracting soils in BaCl2 (Hendershot and Duquette, 1986) and analyzing extracts for Al, Ca, Fe, K, Mg, Mn, and Na cations on an Avio 200 inductively coupled plasma atomic emission spectrometer (Perkin-Elmer, Waltham, Massachusetts, USA). Cation exchange capacity was calculated based on the sum of total cation cmol + charge per kg. The % base saturation was calculated by dividing the sum of base cation concentrations (Ca, K, Mg, and Na) by the total CEC.

### Extracellular enzyme activities

To assess soil microbial activity and potential biogeochemical process rates, we measured the activity of C-degrading, N-degrading, and P-degrading hydrolytic and oxidative enzymes (Table S1) as described in Saiya-Cork et al. (2002) and Hobbie et al. (2012). Briefly, ∼0.5 g soil subsamples were homogenized in a blender with a maleate buffer (100 mM, pH 6.5, based on approximate average pH of soils in our study) and dispensed into 96 well plates for hydrolytic enzymes and 1.5 mL posiclick tubes for oxidative enzymes. Samples were incubated at 20° C in the dark with blanks, standards, and controls for ∼3.5 hours for hydrolytic and ∼20 hours for oxidative assays. Linearity in the change in fluorescence and absorbance over these time frames was confirmed with prior testing. The potential activities of hydrolytic enzymes, α-Glucosidase (AG), β-Glucosidase (BG), Cellobiohydrolase (CBH), β-xylosidase (BX), N-acetyl-β-D-glucosaminidase (NAG), and Acid phosphatase (AP), were determined fluorometrically on a SpectraMaxM2 microplate spectrophotometer (460 nm; Molecular Devices, San Jose, California, USA) using methylumbelliferone (MUB)-labeled substrates (1000 μM for AG, BG, CBH, and NAG, and 2000 μM for BX and AP, excitation at 365 nm, emission at 450 nm). The potential activities of oxidative enzymes, phenol oxidase (PO) and peroxidase (PX), were determined based on absorbance on the SpectraMaxM2 using L-3,4-dihydroxyphenylalanine (L-DOPA) or L-DOPA and hydrogen peroxide as substrates, respectively. Extracellular enzyme activity was calculated from the linear change in fluorescence (hydrolytic enzymes) and absorbance (oxidative enzymes) after correcting for controls, blanks, and soil interference. Cumulative C-degrading hydrolytic EEAs were calculated based on the sum of AG, BG, CBH, and BX while cumulative oxidative EEAs were calculated based on the sum of PO and OX.

### Microbiome analysis

To determine whether microbial communities (general fungi, EM fungi, and bacteria/archaea) differed among EM tree genera, we used high-throughput DNA sequencing. DNA was extracted from 500 mg of freeze-dried soil using the FastDNA SPIN Kit for Soils (MP Biomedicals, Santa Ana, USA). The extracts were purified using cetyl trimethyl ammonium bromide (CTAB). DNA extracts were submitted to the Roy J. Carver Biotechnology Center at the University of Illinois at Urbana-Champaign for Fluidigm amplification (Fluidigm, San Francisco, USA) and Illumina sequencing (Illumina, San Diego, USA). We assessed fungal communities via the *ITS2* gene region using ITS3 and ITS4 (White et al., 1990) primers to amplify DNA. We assessed bacterial and archaeal communities via the bacterial and archaeal *16S* rRNA genes, which were amplified using V4_515F forward (Parada et al., 2016) and V4_806R reverse primers (Apprill et al., 2015). Fungal and bacterial/archaeal amplicons were sequenced via NovaSeq 2 x 250 bp. Primer sequences and references can be found in Table S2. Sequence data is publicly available from the NCBI SRA database under accession number (to be provided upon manuscript acceptance).

We used the DADA2 pipeline (Callahan et al., 2016) for bioinformatic processing to produce amplicon sequence variants (ASVs) from the sequencing data. Briefly, quality filtering, denoising, merging forward and reverse reads, and removing chimeric sequences was performed using recommended parameters for *16S* and *ITS* genes (Callahan et al., 2016). We did not cluster sequence variants (Glassman and Martiny, 2018) prior to using the default DADA2 classifier (Wang et al., 2007) to assign taxonomy based on reference sequences from the UNITE (v10.0) database (Kõljalg et al., 2013) for *ITS* sequences and the SILVA (R138.1) database (Quast et al., 2012) for *16S* sequences. Data used for analysis consisted of 6991 fungal, 64266 bacterial, and 549 archaeal ASVs. Due to the relatively low number of archaeal sequence variants, and because bacterial and archaeal sequences were derived from the same primer sets, we analyzed bacterial and archaeal communities together. Ectomycorrhizal fungi were assigned at the genus level using the FungalTraits (Põlme et al., 2020) and FUNGuild database (Nguyen et al., 2016) on ASVs designated with a “probable” or “highly probable” confidence score for EM classifications, as well as family level for all Inocybaceae (Matheny et al., 2020). Alpha diversity of EM fungal, general fungal, and bacterial groups was assessed using Hill number at *q* = 1 (analagous to Shannon index; Chao et al., 2014) with the hillR package in R (Li, 2018). To assess microbial taxa with differential abundance among EM tree genera, ASVs were aggregated to genus or higher-level assignments based on the lowest phylogenetic level assigned from the respective taxonomic database.

### Statistical analysis

All statistical analyses were performed in the R statistical environment (R Core Development Team 2013). First, we assessed univariate relationships within and among leaf litter and SOM properties using linear models with tree genus as a fixed effect in the *nlme* package (Pinheiro et al., 2017). We tested for spatial autocorrelation of residuals for variables of interest using Moran’s I in the *ape* package (Paradis et al., 2019). For variables with significant spatial autocorrelation, we incorporated a Gaussian correlation structure based on the X and Y coordinates of focal trees into the linear models. We used this same approach to assess differences in EEA rates and stoichiometries, as well as differences in the relative abundance of microbial taxa and fungal functional guilds. P values from univariate tests were adjusted for multiple comparisons using the BH false discovery rate (Benjamini, 2010, Benjamini and Hochberg, 1995). For tests deemed significant at p < 0.05, we used a post-hoc Tukey HSD test to determine mean separation among tree genera.

We assessed differences among tree genera in SOM biogeochemical variables as well as general fungal community and bacterial/archaeal community composition via distance-based redundancy analysis (dbRDA) with the vegan package in R (Oksanen et al., 2010). We standardized SOM variables using a Z-score transformation and relativized ASV count numbers using a Hellinger transformation to account for differences in overall read numbers among samples (Legendre and Gallagher, 2001). We then created distance matrixes for each variable group using Euclidean distance for SOM variables and Jaccard distance for microbial groups.

These distance matrixes were then used in a dbRDA model with tree genus as a fixed factor and conditioned with physical proximity (X and Y coordinates) to account for spatial autocorrelation, with subsequent ANOVAs permuted 9999 times to calculate p-values. We visually depict these relationships with ordinations using the first and second unconstrained multidimensional scaling (MDS) axis. To describe relationships between microbial community dissimilarity and EM-associated functional traits, we also conducted dbRDAs on fungal and bacterial/ archaeal dissimilarity matrixes with leaf litter C : N, MAOM δ^13^C, litter δ^15^N, EM fungal diversity, oxidative EEA, and the SOM index (MDS1 from SOM ordination) as fixed factors and conditioned on X and Y sampling coordinates. We then ran model selection on these full dbRDAs using the *ordistep* function, and included all factors retained in final ordinations. Statistical significance was assessed as p < 0.05 unless otherwise stated.

## Results

In support of our hypothesis, we found tree genera differed across functional traits related to litter quality, resource use patterns, and microbiome assembly (Figure 2). Functional trait differences among EM tree genera also generally correlated with SOM properties in surrounding soils. Overall, *Quercus* had significantly higher leaf litter C : N, EM fungal diversity, oxidative EEA and lower MAOM δ^13^C and litter δ^15^N than *Tilia* (p < 0.05), with *Carya* frequently falling intermediary.

**Figure 2.**
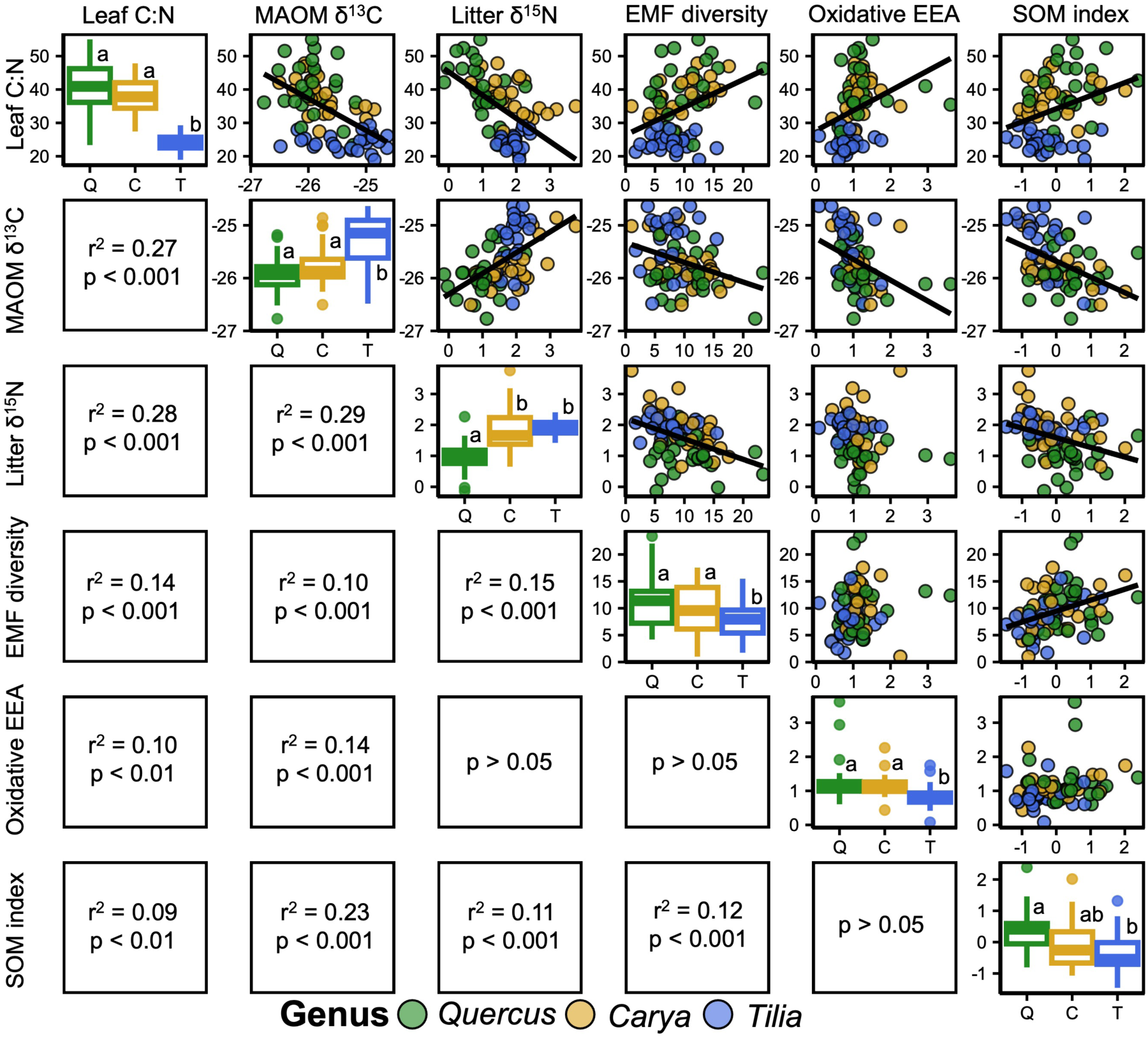
Differences among tree genera for leaf litter C : N, MAOM δ^13^C, leaf litter δ^15^N, ectomycorrhizal fungal (EMF) diversity as Hill # at *q* = 1 (Shannon index), cumulative oxidative extracellular enzyme activity (EEAs: nM substrate g soil^-1^ hour^-1^; diagonal), and the soil organic matter (SOM) index expressed as the first multidimensional scaling axis from the principial coordinates analysis shown in figure 3. Letters in the diagonal plots represent groups that significantly differ from one another at p < 0.05. Intersections depict correlations among these variables above the diagonal and results from corresponding linear regression below the diagonal.

Ectomycorrhizal tree genera differed in C and N isotopic composition of leaf litter and SOM, potentially corresponding to alternative resource acquisition and allocation patterns. In leaf litter, *Quercus* had higher δ^13^C and lower δ^15^N than *Carya* and *Tilia* while in SOM fractions δ^13^C and δ^15^N were lowest near *Quercus* trees (Figure 3AB). These differences led to alternative patterns of differences in C and N isotopic values between leaf litter and soil for *Quercus* trees than the other two genera, with lower δ^13^CMAOM-litter and higher δ^15^NMAOM-litter than *Carya* and *Tilia* (Figure 3C). The relative strength of litter to SOM isotopic difference was correlated for C and N such that smaller differences in δ^13^C were associated with smaller differences in δ^15^N (Figure 3D; r^2^ = 0.21, p < 0.001).

**Figure 3.**
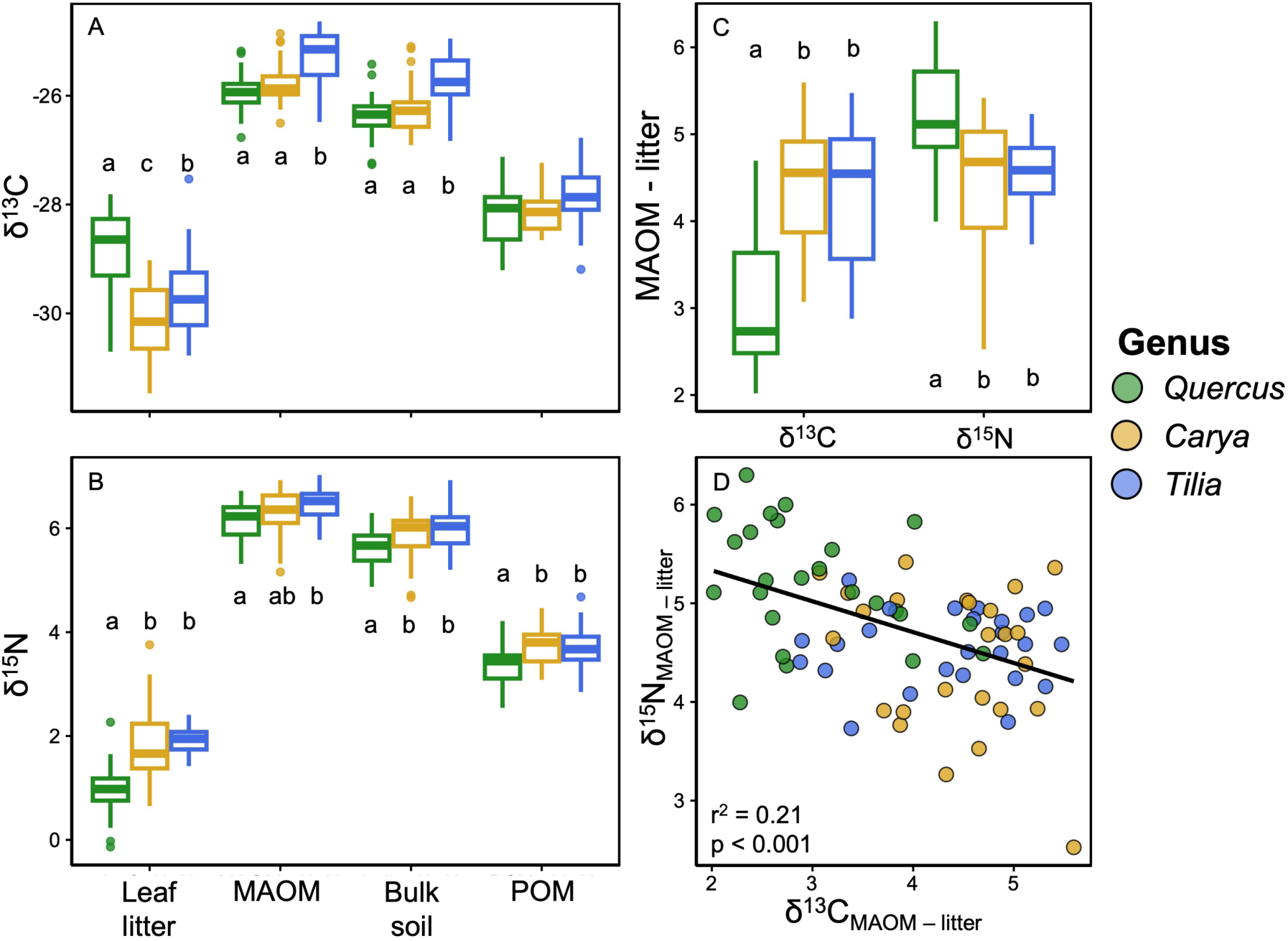
Values of δ^13^C (A) and δ^15^N (B) in leaf litter, mineral-associated organic matter (MAOM), bulk soil, and particulate organic matter (POM) for each tree genus. Differences between the MAOM and leaf litter δ^13^C and δ^15^N values (C) and the correlation between these differences (D) could represent fractionating forces related to resource use and allocation among tree genera. Letters represent groups significantly differing from one another (p < 0.05) within leaf litter and SOM fractions (AB) and isotopic difference (C).

Soil organic matter properties differed among EM tree genera, primarily among C and N dynamics (Figure 4, Table 1). Overall, EM tree genera explained 6% of variation in SOM dissimilarity (p = 0.003), after accounting for spatial auto correlation. Soil C and N concentrations and properties significantly differed among tree genera, with *Tilia* having the lowest bulk soil %C and %N, dissolved organic C, microbial biomass C, and dissolved N (p < 0.05; Table 1). Yet, there were no differences among genera for soil inorganic N (NH^+^ and NO3^-^) concentrations (p > 0.05, Table 1). These C and N SOM properties were scaled along MDS1 of the SOM ordination, which explained 23% of overall variation in SOM properties and differed significantly between *Quercus* and *Tilia* trees (p < 0.01, Figure 4). The second SOM ordination axis correlated with general measures of acid-base chemistry (*i.e.,* pH, CEC, or % base saturation) and did not differ among EM tree genera (Figure 4; Table 1).

**Figure 4.**
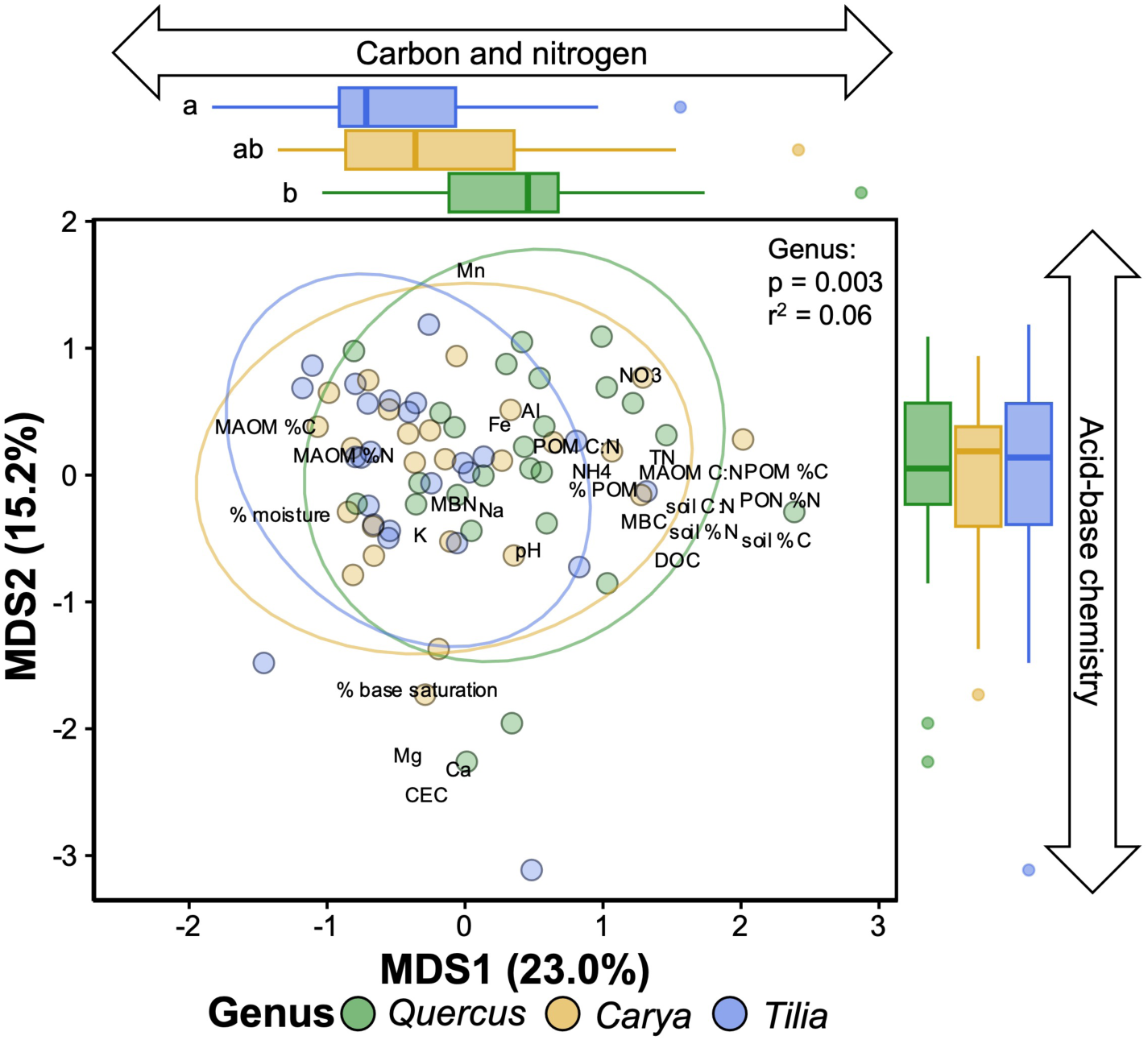
Ordination for constrained principal correspondence analysis of soil organic matter (SOM) variables conditioned in sampling coordinates to remove the influence of spatial autocorrelation. Axes are labeled with the proportion of variation explained by the first (x) and second (y) multi-dimensional scaling (MDS) component. Labels are the endpoint vector loadings of each SOM variable used to construct the ordination, multiplied by two to increase visibility (Table 1). Boxplots of genera scores for MDS1 and MDS2 are shown in the margins, with letters indicating significant differences at p < 0.05.

**Table 1.** Differences among EM tree genera in SOM biogeochemical variables. P-values were adjusted for multiple comparisons using the false discover rate. Bolded values are significant at p < 0.05. Values under species labels represent mean ± standard error, values with different letters are significantly different from one another at p < 0.05.

| Variable | Moran's I | F-value | p-value adjusted | *Quercus* | *Carya* | *Tilia* |
| --- | --- | --- | --- | --- | --- | --- |
| pH | 0.141 | 0.223 | 0.838 | 6.58 ± 0.04 | 6.62 ± 0.04 | 6.59 ± 0.03 |
| % water content | 0.535 | 3.289 | 0.079 | 0.59 ± 0.01 | 0.59 ± 0.01 | 0.62 ± 0.01 |
| **DOC** | **0.060** | **20.038** | **<0.001** | 41.25 ± 2.21 (a) | 27.7 ± 2.01 (b) | 24.53 ± 1.67 (b) |
| **TDN** | **0.282** | **7.896** | **0.003** | 12.89 ± 0.7 (a) | 9.85 ± 0.63 (b) | 9.78 ± 0.54 (b) |
| **MBC** | **<0.001** | **6.974** | **0.005** | 366.71 ± 16.66 (a) | 338.57 ± 20.89 (a) | 276.34 ± 14.32 (b) |
| MBN | 0.102 | 2.243 | 0.150 | 4.60 ± 0.23 | 5.43 ± 0.37 | 4.64 ± 0.31 |
| NH<sub>4</sub><sup>+</sup> | 0.298 | 0.649 | 0.598 | 2.35 ± 0.11 | 2.12 ± 0.13 | 2.05 ± 0.28 |
| NO<sub>3</sub><sup>-</sup> | <0.001 | 2.685 | 0.113 | 7.61 ± 0.72 | 5.90 ± 0.47 | 6.10 ± 0.47 |
| Total % MAOM | 0.047 | 2.837 | 0.102 | 87.72 ± 0.01 | 87.62 ± 0.01 | 90.43 ± 0.01 |
| **MAOM $\delta^{13}\text{C}$** | **<0.001** | **12.440** | **<0.001** | -25.93 ± 0.07 (a) | -25.77 ± 0.07 (a) | -25.32 ± 0.1 (b) |
| MAOM %C | 0.946 | 2.113 | 0.177 | 0.94 ± 0.02 | 0.95 ± 0.01 | 0.99 ± 0.01 |
| **MAOM $\delta^{15}\text{N}$** | **<0.001** | **4.523** | **0.031** | 6.15 ± 0.07 (a) | 6.32 ± 0.09 (ab) | 6.48 ± 0.06 (b) |
| MAOM %N | 0.397 | 0.156 | 0.838 | 1.00 ± 0.02 | 1.01 ± 0.01 | 1.02 ± 0.02 |
| MAOM C:N | 0.829 | 0.980 | 0.596 | 12.20 ± 0.11 | 12.05 ± 0.16 | 11.94 ± 0.1 |
| POM $\delta^{13}\text{C}$ | <0.001 | 3.462 | 0.065 | -28.2 ± 0.1 | -28.08 ± 0.08 | -27.83 ± 0.11 |
| POM %C | 0.606 | 1.757 | 0.249 | 0.19 ± 0.01 | 0.16 ± 0.01 | 0.15 ± 0.01 |
| **POM $\delta^{15}\text{N}$** | **0.026** | **6.892** | **0.035** | 3.36 ± 0.08 (a) | 3.75 ± 0.07 (b) | 3.69 ± 0.07 (b) |
| POM %N | 0.752 | 1.323 | 0.357 | 0.12 ± 0.01 | 0.10 ± 0.01 | 0.10 ± 0.01 |
| POM C:N | 0.885 | 1.003 | 0.357 | 20.48 ± 0.43 | 19.97 ± 0.74 | 19.33 ± 0.50 |
| **Bulk soil $\delta^{13}\text{C}$** | **<0.001** | **11.518** | **<0.001** | -26.40 ± 0.08 (a) | -26.21 ± 0.09 (a) | -25.74 ± 0.11 (b) |
| **Bulk soil %C** | **0.010** | **7.326** | **0.004** | 8.54 ± 0.22 (a) | 8.05 ± 0.33 (a) | 7.09 ± 0.23 (b) |
| **Bulk soil $\delta^{15}\text{N}$** | **<0.001** | **5.348** | **0.017** | 5.60 ± 0.07 (a) | 5.86 ± 0.09 (b) | 5.99 ± 0.07 (b) |
| **Bulk soil %N** | **0.001** | **5.893** | **0.012** | 0.65 ± 0.01 (a) | 0.62 ± 0.02 (ab) | 0.56 ± 0.01 (b) |
| Bulk soil C:N | 0.460 | 3.060 | 0.092 | 12.98 ± 0.15 | 12.87 ± 0.19 | 12.46 ± 0.1 |
| **Leaf $\delta^{13}\text{C}$** | **0.902** | **15.387** | **<0.001** | -28.95 ± 0.17 (a) | -30.18 ± 0.13 (c) | -29.65 ± 0.15 (b) |
| **Leaf %C** | **0.003** | **112.371** | **<0.001** | 49.38 ± 0.17 (a) | 46.20 ± 0.16 (b) | 44.52 ± 0.32 (c) |
| **Leaf $\delta^{15}\text{N}$** | **<0.001** | **25.097** | **<0.001** | 0.95 ± 0.1 (a) | 1.86 ± 0.14 (b) | 1.93 ± 0.05 (b) |
| **Leaf %N** | **<0.001** | **59.741** | **<0.001** | 1.26 ± 0.05 (a) | 1.24 ± 0.03 (a) | 1.86 ± 0.04 (b) |
| **Leaf C:N** | **<0.001** | **60.641** | **<0.001** | 40.63 ± 1.61 (a) | 37.92 ± 1.01 (a) | 24.14 ± 0.49 (b) |
| CEC | 0.667 | 0.102 | 0.799 | 55.05 ± 2.41 | 55.58 ± 1.94 | 54.15 ± 2.38 |
| % base saturation | 0.681 | 0.154 | 0.799 | 99.64 ± 0.07 | 99.65 ± 0.05 | 99.58 ± 0.12 |

Oxidative EEAs differed among EM tree genera (p < 0.05), but hydrolytic EEAs did not. Cumulative activities of oxidative enzymes were lower under the least nutrient conservative *Tilia* than the more nutrient conservative *Quercus* and *Carya* species (p < 0.05, Figure 2). Activities of peroxidase and phenol oxidase considered individually differed only between *Quercus* and *Tilia*, with *Carya* not differing significantly from either (Table S3). Generally, there were no significant differences among tree genera for hydrolytic EEAs, though β-xylosidase activity was higher under *Quercus* than *Tilia* (p < 0.05, Table S3). The ratio of cumulative oxidative : cumulative hydrolytic C degrading EEAs was also marginally greater under *Quercus* and *Carya* than *Tilia* (p = 0.07, Table S3).

Microbial community composition differed among EM tree genera, but fungal communities had stronger correlations with EM tree functional traits than bacterial/ archaeal. Tree genus explained ∼3% of variation in fungal and bacterial/ archaeal communities after accounting for spatial autocorrelation (p < 0.01, Figure 5AB). Fungal community composition significantly corresponded to functional traits related to resource use (MAOM δ^13^C and leaf δ^15^N), as well as EM fungal diversity and SOM properties (p < 0.05; Figure 5C), while bacterial/ archaeal community composition was related with MAOM δ^13^C and EM fungal diversity (p < 0.05; Figure 5D). Neither microbial group were significantly associated with leaf litter quality and bacterial/ archaeal composition did not correlate with the SOM index (p > 0.05). Models accounting for functional traits explained over double the variation of tree genus in fungal community composition (7% vs 3%) while these models explained similar variation in bacterial/ archaeal communities (∼3%).

**Figure 5.**
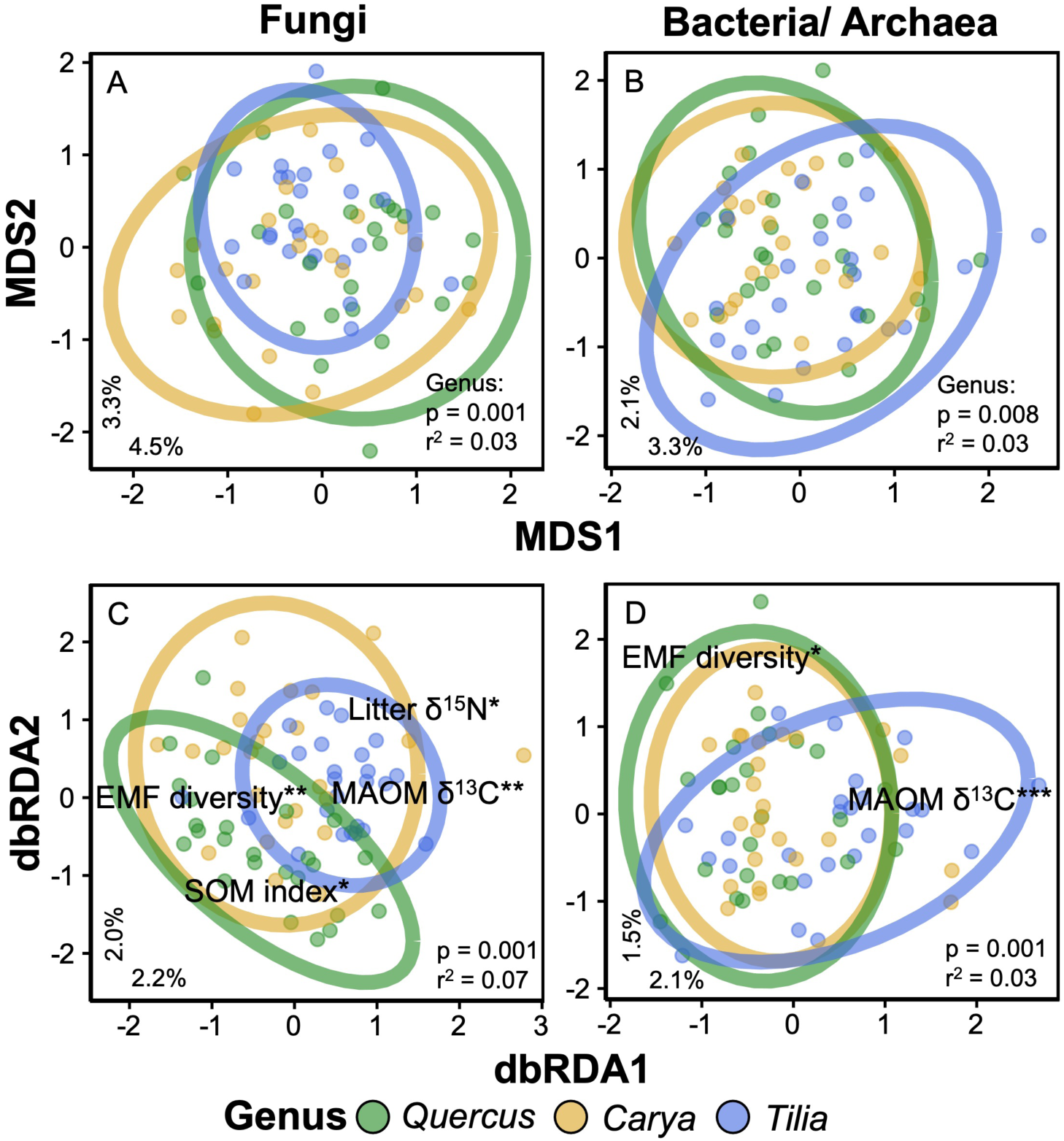
Ordinations for constrained principal coordinates analysis (AB) and distanced- based redundancy analysis (dbRDA, CD) of fungal and bacterial/ archaeal communities. All ordinations were conditioned based on sampling coordinates to remove the influence of spatial autocorrelation. Ellipses represent 95% confidence intervals about the centroid for each genus. The proportion of variation explained by the first (x) and second (y) multi- dimensional scaling (MDS) or dbRDA axis are shown in the lower left corner of each plot. Variable labels in panels C and D represent the vector endpoints of functional trait variables (Figure 3) that significantly correlated with community dissimilarity based on model selection (*p<0.1, **p<0.01, ***p<0.001).

The diversity of microbial groups and relative abundance of specific microbial taxa differed among tree genera. The alpha diversity of both fungi overall and EM fungi specifically was greatest under *Quercus* trees and lowest under *Tilia* (p > 0.05), yet the diversity of bacteria and the relative abundance of fungal functional guilds did not differ among genera (Figure S2). When taxonomic assignments were aggregated into genus or higher-level identifications, *Inocybe* (EM) was the most abundant taxon across all three tree genera. *Inocybe* comprised roughly 23% of the fungal community under *Carya,* 15% under *Quercus,* and 11% under *Tilia*, and was significantly greater under *Carya* than *Tilia* (F2,72 = 3.78, p < 0.05; Figure S3). None of the other dominant fungal taxa (*Mortierella,* Agaricales, *Preussia, Tuber,* and *Tausonia*) differed in abundance among tree genera (p > 0.05, Figure S3). Several dominant bacterial taxa were more abundant under *Tilia* than other tree genera (Gaiellales and Xanthobacteraceae; p < 0.05, Figure S4), while others (Vicinamibacteriales, Nitrososperaceae, Gaiella, and Udaeobacter) did not differ among tree genera (p > 0.05, Figure S4).

## Discussion

While EM trees are often generalized as a unified functional guild in mycorrhizal paradigms (Talbot et al., 2015), our study shows that broadleaf EM tree genera can exhibit divergent functional traits that correspond to alternative ecosystem functions. In support of our hypothesis, there were consistent differences between *Quercus* and *Tilia* trees related to their litter quality, resource acquisition and allocation patterns, and microbiome assembly/ function. These functional differences correlated with SOM dynamics, which could be indicative of broader variation in ecosystem response to EM tree abundance. The differences we found among these EM tree genera demonstrate the potential importance of considering the interactions between ecophysiology and biodiversity in assessing broad-scale ecosystem patterns (Lavorel and Garnier, 2002). These patterns may represent an emergent axis of functional diversity among EM mutualisms and its influence on ecosystem function. Overall, our findings show that EM mutualisms can display significant functional diversity, suggesting that EM association should not be considered a homogenous trait of tree species in predicting their effects on ecosystems.

The EM tree genera we sampled within a single forest stand differed in patterns of resource allocation and acquisition. We found differences among EM tree genera, particularly *Quercus* and *Tilia*, in leaf litter nutrient stoichiometry, SOM δ^13^C, and leaf litter δ^15^N, as well as correlations among these variables potentially representing a spectrum of nutrient conservative and mycorrhizal-supportive traits. *Quercus* was generally on the more conservative/ collaborative end of this spectrum, with higher leaf litter C : N, smaller differences in C and larger differences in N isotopic compositions between leaf litter and SOM, while *Tilia* showed the opposite pattern. *Carya* was somewhat intermediate, displaying conservative nutrient strategies but more limited collaborative resource transfer. Broadly, there is significant interspecific variation in leaf litter quality across EM tree taxa (Sun et al., 2018) and in belowground C allocation in temperate forests with varying EM tree communities (Keller et al., 2021). Ecological trade-offs favoring various economic strategies are common (Reich, 2014); thus it is likely that trees investing more C belowground to acquire nutrients from mycorrhizal fungi may also conserve these nutrient in their foliar tissues (Jiang et al., 2023). More conservative nutrient-use strategies are commonly associated with EM tree species (Averill et al., 2019), though this is often attributed to their greater relative abundance at sites with lower soil fertility (Mao et al., 2019). However, we did not find differences within our study site in concentrations of soil inorganic N or measures of soil acid-base chemistry, a potential mediator of plant/ soil isotopic composition and mycorrhizal resource economics (Lin et al., 2021, Seyfried et al., 2022). The persistence of these resource strategies independent of environmental homogeneity indicates that they may be conserved at the genus level.

Mycorrhizal C and N exchange is often considered linear, with greater C allocation resulting in greater N acquisition (Bogar et al., 2022, Horning et al., 2023); however, this relationship is not universal (Hasselquist et al., 2016, Näsholm et al., 2013, Plett et al., 2020) and may not be directly proportional at all (Bunn et al., 2024). We found significant correlations between derived measures of plant belowground C allocation (δ^13^CMAOM-litter) and mycorrhizal N acquisition (δ^15^NMAOM-litter), supporting the reciprocal transfer of these resources. In arbuscular mycorrhizal symbiosis, a plant host can directly mediate these trades, receiving more or less N per unit of C supplied (Fellbaum et al., 2014). However, trade-balance models suggest that the availability and allocation of resources should covary based on their respective need (Johnson, 2010), yet these frameworks insufficiently explain C reciprocation for nutrient acquisition (Corrêa et al., 2023). Alternatively, source-sink dynamics, where plant resource allocation is based on both its individual need and the soil availability may be more appropriate for EM mutualisms (Bogar, 2023). These differences in the C cost of mycorrhizal N acquisition may also be influenced by transfers / availability of other nutrients or non-nutritional benefits provided by EM fungi (Hortal et al., 2017, Berrios et al., 2023). Source-sink dynamics would suggest that trees with a C surplus should allocate this resource belowground due to inherent nutrient limitation (Bunn et al., 2024), regardless of reciprocal exchange. However, this pattern would still support the relationship observed here, where trees with greater C surplus from nutrient limitation also have greater reliance on their mycorrhizal mutualists for nutrient acquisition to combat this limitation. Many current resource models to understand mycorrhizal resource transfer do not fully account for the role of EM fungi in driving ecosystem biogeochemical syndromes (Phillips et al., 2013) to potentially exacerbate or reinforce edaphic nutrient limitations, or coevolution for specific functions between partners (Hoeksema, 2010). Here, we show that EM tree genera may differ in the strength of their mycorrhizal relationship relative to C and N resource transfer within a given forest stand, which could also be tied to EM-associated impacts on surrounding soil ecosystems.

Leaf litter quality may be related to patterns in belowground resource transfer. Both leaf and root litter influence the formation and turnover of SOM (Hobbie, 2015), but leaf litter traits are important determinants of overall decomposition and nutrient cycling in ecosystems (Mooshammer et al., 2012, Rahman et al., 2013). Lower quality leaf litter increases a tree’s reliance on EM fungi for nutrient acquisition (Trap et al., 2017). While differences in leaf litter quality among these genera has been reported previously (Craig et al., 2018), there is less evidence for their connection with mycorrhizal resource transfer. *Tilia* saplings can allocate marginally less C to belowground root tissues than *Quercus* saplings (Keller and Phillips, 2019). Mature *Tilia* also demonstrate faster rates of mineral N uptake than *Quercus*, while *Quercus* prefer organic N (Schulz et al., 2011). A greater C cost and reliance on mycorrhizal fungi for N acquisition for *Quercus* would indicate a higher resource value, which could lead to more nutrient conservative strategies for this genus (Fisher et al., 2010). Higher leaf litter C : N ratio in *Quercus* and *Carya* species may have been promoted by higher C allocation to leaf defensive structures (i.e., lignin or tannins; Mason and Donovan, 2015), or greater reabsorption of N prior to leaf senescence (Zhang et al., 2018). Higher quality litter increases N mineralization rates (Scott and Binkley, 1997, Fanin and Bertrand, 2016), promoting availability of inorganic N. Indeed, *Tilia* can have higher nutrient turnover rates from litter than *Quercus* (Schmidt et al., 2016), supporting their lower reliance on EM fungi for nutrient acquisition. Root traits may also align with these patterns, as faster decomposing fine roots are also associated with lower C : nutrient stoichiometries in root tissues (Carrillo et al., 2017, Gao et al., 2021). Our results show that leaf litter quality can override general effects of EM association in their relationship with SOM dynamics. It may be important to account for this interspecific functional trait variation in predicting the effect of EM trees on ecosystem function.

We found that microbial communities for general fungi and bacteria/ archaea differed among EM tree genera and consistently corresponded to EM tree functional traits.

Rhizodeposition is one of the strongest tools for plants to influence microbiome assembly (Ladygina and Hedlund, 2010, Paterson et al., 2007), with nutrient trading often mediating the magnitude of this flux (Jones et al., 2009). Rhizosphere microbiomes, particularly EM fungi, supplied with more plant photosynthate C would have greater capacity to liberate nutrients from SOM via extracellular enzymes (Kaiser et al., 2010). Activities of oxidative enzymes necessary for degradation of complex SOM, like lignin, can be up to 90% greater in presence of plant roots relative to adjacent trenched plots (Sterkenburg et al., 2018). Supporting the strong relationship we found between rates of oxidative EEAs and both leaf litter chemical quality and MAOM δ^13^C. These patterns may also be reflected in the diversity of EM fungi. Ectomycorrhizal host specificity can vary widely between plant and fungal species (Peay et al., 2015, Tedersoo et al., 2008), with biogeographic or environmental factors also influencing this complementarity (Li et al., 2023). EM fungal alpha diversity was higher under *Quercus* and *Carya* than *Tilia*, potentially due to mycorrhizal support, as greater belowground C allocation would create more niche space for EM fungi. Similar patterns can be found for other belowground functional traits, like root morphology (Li et al., 2017). Overall, EM trees appear to assemble distinct soil microbiomes that may be related to emergent properties in the influence on ecosystem function.

Interspecific variation among EM trees in their functional traits could be important for using mycorrhizal associations to predict ecosystem function and response to global change. The relationship between the relative abundance of EM trees and soil C dynamics is not consistent at larger spatial scales (Lin et al., 2017, Soudzilovskaia et al., 2019), which also likely reflects compositional dissimilarity among EM tree communities. As soil C is generally more stable when derived from belowground inputs rather than leaf litter (Sokol and Bradford, 2019), differential trends in root C allocation and its influence on SOM among EM tree species may contribute to highly variable soil C stocks in EM-dominated ecosystems. Further, leaf litter traits can exert strong controls on microbial turnover (Craig et al., 2022) and C use efficiency (Ridgeway et al., 2022) that also likely influence the ultimate stabilization and accumulation of soil C (Tao et al., 2023). Broadly, EM-associating plant species are expected to respond to increasing atmospheric CO2 concentrations with greater investments in plant biomass, rather than soil C (Terrer et al., 2021). However, if these plant species differ in their tissue-soil C allocation strategies, then this would also likely affect how EM-dominated ecosystems with differing species assemblages would respond to global change. Reliance on organic soil N acquitted from EM fungi has been proposed to negatively affect the capacity of EM dominated ecosystems to respond to increasing anthropogenic N deposition (Averill et al., 2018). Our evidence suggests that some EM tree species may be able to respond to these shifting nutrient conditions better than others, potentially complicating predictions of overall EM ecosystem decline in response to global change (Jo et al., 2019). There may also be plasticity in EM tree species’ reliance on mycorrhizal mutualists based on environmental conditions (Pellitier et al., 2021), indicating a need for further exploration into the dynamic relationship between EM biodiversity and resource economics.

## Conclusion

Overall, we show that there is significant functional variation among EM-associating tree genera that can directly mediate surrounding ecosystem function and SOM dynamics. This relationship could potentially promote EM-associated biogeochemical syndromes for some EM trees, enriching SOM-C stocks and altering nutrient cycling dynamics via low quality litter inputs and increase EM fungal activity. However, EM trees with higher litter quality can have alternative resource use and microbiome assembly patterns that may drive ecosystem outcomes that do not align with conventional mycorrhizal paradigms. This unaccounted variation among EM-associating tree species may be responsible for the inconsistencies of general EM effects on ecosystem function. Properly accounting for phylogenetic and other functional variation in these EM traits could provide more accurate models of mycorrhizal SOM properties. We demonstrate that EM mutualisms are not functionally homogenous among broadleaf tree genera, with potential implications for the generalizability of coarse mycorrhizal associations in understanding forest ecosystem function.

## Author contribution

JDE and WHY conceived of the ideas and designed the methodology; JDE and WCE collected the data; JDE analyzed the data and led the writing of the manuscript; JWD and JMF maintained the sampling site; All authors contributed critically to the drafts and gave final approval for publication.

## Data availability

Data and analytical code for this project are publicly available at https://github.com/jedward-s/2024EctoResource. Sequence data is publicly available from NCBI under SRA #(to be provided upon manuscript acceptance).

## Acknowledgements

We would like to thank Ebony Rankin, Sydney Bell, Chuck Hyde, Sean Khan Ooi, Rachel van Allen, Adam von Haden, and Georgia Seyfried for their help collecting and analyzing the data in this study. We would also like to thank Angela Kent, Anthony Yannarell, Adam Davis, and Sören Weber for providing comments of the study design, analysis, and previous versions of this manuscript. JDE was supported by the Cooperative State Research, Education, and Extension Service, US Department of Agriculture, under project number ILLU 875-952, as well as by NSF DEB award #2305863.

