## Supplemental for "Functional traits of ectomycorrhizal trees influence their effects on surrounding soil organic matter properties"

**Table S1.** Names, types, and putative function of extracellular enzymes assayed in this study.

| **Enzyme** | **Type** | **Putative function** |
|---|---|---|
| α-Glucosidase (AG) | Hydrolytic | Degrades starch |
| β-Glucosidase (BG) | Hydrolytic | Degrades cellulose |
| Cellobiohydrolase (CBH) | Hydrolytic | Degrades cellulose |
| β-xylosidase (BX) | Hydrolytic | Degrades hemicellulose |
| N-acetyl-β-D-glucosaminidase (NAG) | Hydrolytic | Degrades organic N |
| Acid phosphatase (AP) | Hydrolytic | Degrades organic P |
| Phenol oxidase (PO) | Oxidative | Degrades lignin (and other SOM) |
| Peroxidase (PE) | Oxidative | Degrades lignin (and other SOM) |

**Table S2.** Target genes, PCR primer names, primer sequences, and references for genes sequences in fungal and bacterial/ archaeal community analysis.

| **Target gene** | **PCR primers** | **Primer sequence (5'→3')** | **Reference** |
|---|---|---|---|
| Fungal ITS | ITS3 | GCATCGATGAAGAACGCAGC | White et al 1990 |
| | ITS4 | TCCTCCGCTTATTGATATGC | |
| Bacterial and archaeal 16S | V4_515F | GTGYCAGCMGCCGCGGTAA | Parada et al. 2016 Apprill et al 2015 |
| | V4_806R | GGACTACNVGGGTWTCTAAT | |

**Table S3.** Differences among tree genera for extracellular enzyme activities determined using general linear models. Units for all EEAs are nM substrate g soil$^{-1}$ hour$^{-1}$. Values in the tree genus columns represent mean ± standard error, with different letters indicating statistically significant differences at $p < 0.05$.

| **Variable** | **F-value** | **p-value adjusted** | ***Quercus*** | ***Carya*** | ***Tilia*** |
|---|---|---|---|---|---|
| AG | 1.413 | 0.250 | 37.48 ± 1.29 | 34.36 ± 1.19 | 39.64 ± 1.38 |
| BG | 1.226 | 0.299 | 510.4 ± 12.1 | 490.2 ± 12.2 | 467.2 ± 9.30 |
| CBH | 1.595 | 0.209 | 180.8 ± 6.88 | 181.3 ± 5.96 | 159.0 ± 4.30 |
| **BX** | **3.253** | **0.044** | 245.2 ± 7.15 (a) | 234.5 ± 7.65 (ab) | 215.5 ± 6.67 (b) |
| Cumulative hydrolytic C degrading enzymes | 1.404 | 0.252 | 842.4 ± 21.2 | 810.8 ± 20.1 | 764.4 ± 15.4 |
| NAG | 1.252 | 0.291 | 113.8 ± 2.70 | 105.0 ± 2.28 | 98.52 ± 2.34 |
| AP | 0.353 | 0.703 | 697.1 ± 25.5 | 601.0 ± 16.94 | 561.4 ± 16.1 |
| **PO** | **5.002** | **0.009** | 656.4 ± 39.4 (a) | 594.8 ± 20.1 (ab) | 445.1 ± 20.0 (b) |
| **PX** | **5.764** | **0.005** | 625.6 ± 38.2 (a) | 556.3 ± 19.8 (ab) | 404.4 ± 19.2 (b) |
| **Cumulative oxidative enzymes** | **5.420** | **0.006** | 1282 ± 77.4 (a) | 1151 ± 39.4 (a) | 849.5 ± 39.1 (b) |
| Ratio of cumulative oxidative enzymes: cumulative hydrolytic C degrading enzymes | 2.705 | 0.073 | 1.52 ± 0.07 | 1.46 ± 0.05 | 1.14 ± 0.05 |

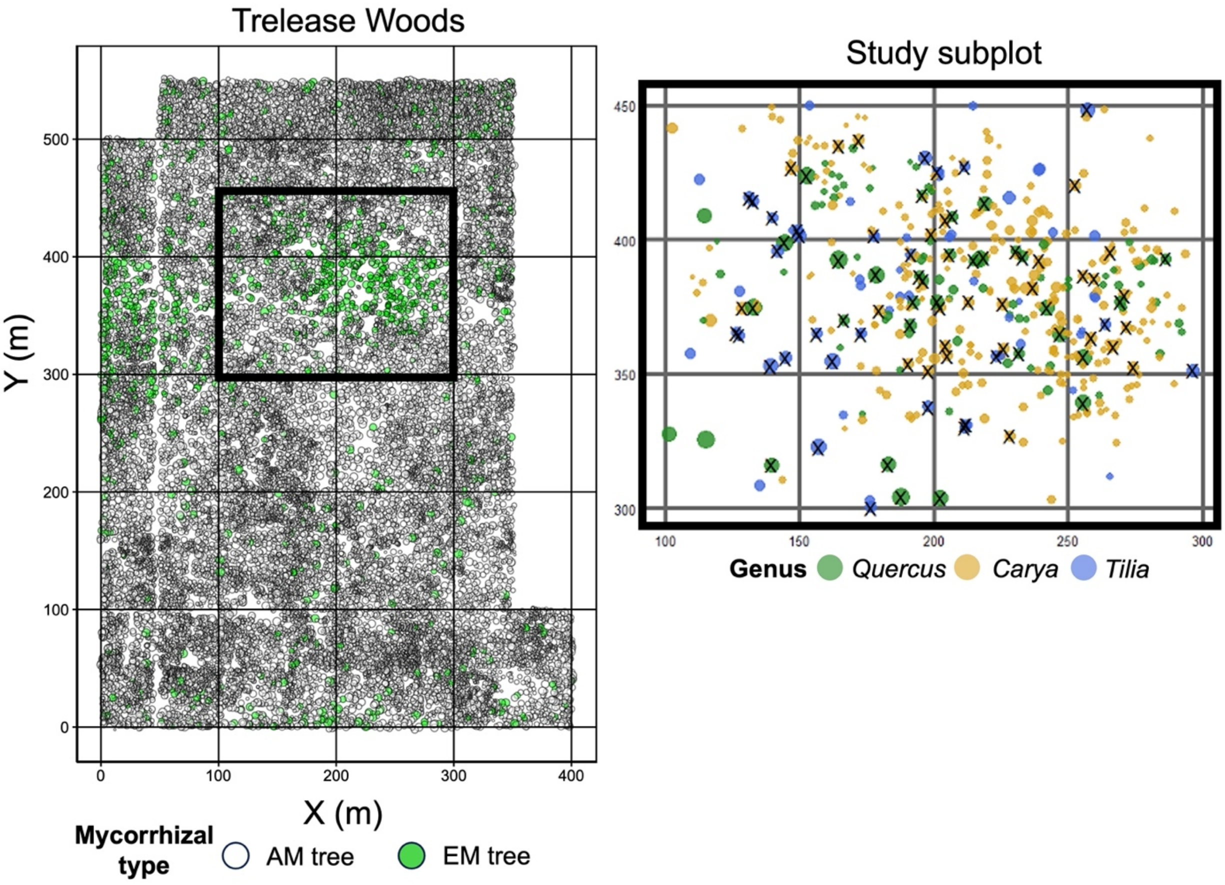


**Figure S1.** Map of mycorrhizal association of all arbuscular mycorrhizal (AM) and ectomycorrhizal (EM) associating trees within Trelease woods (left) and genera of EM-association trees within study subplot (right). Point size is proportional to tree DBH, Xs indicate a focal tree sampled in this study.

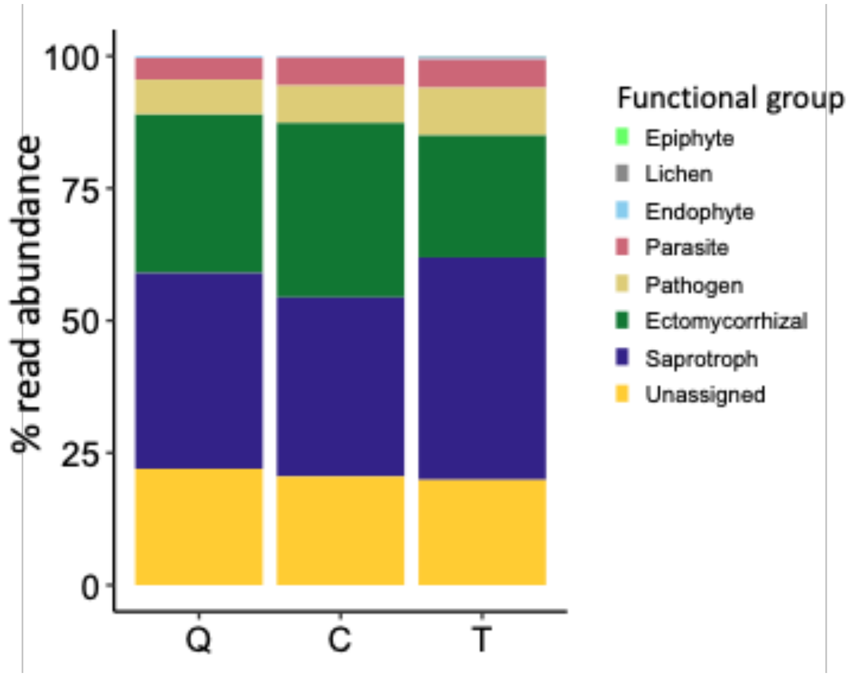


Figure S2. Functional group composition of fungal communities from *Quercus* (Q), *Carya* (C), and *Tilia* (T) trees based on assignments from FungalTraits and FUNGuild.

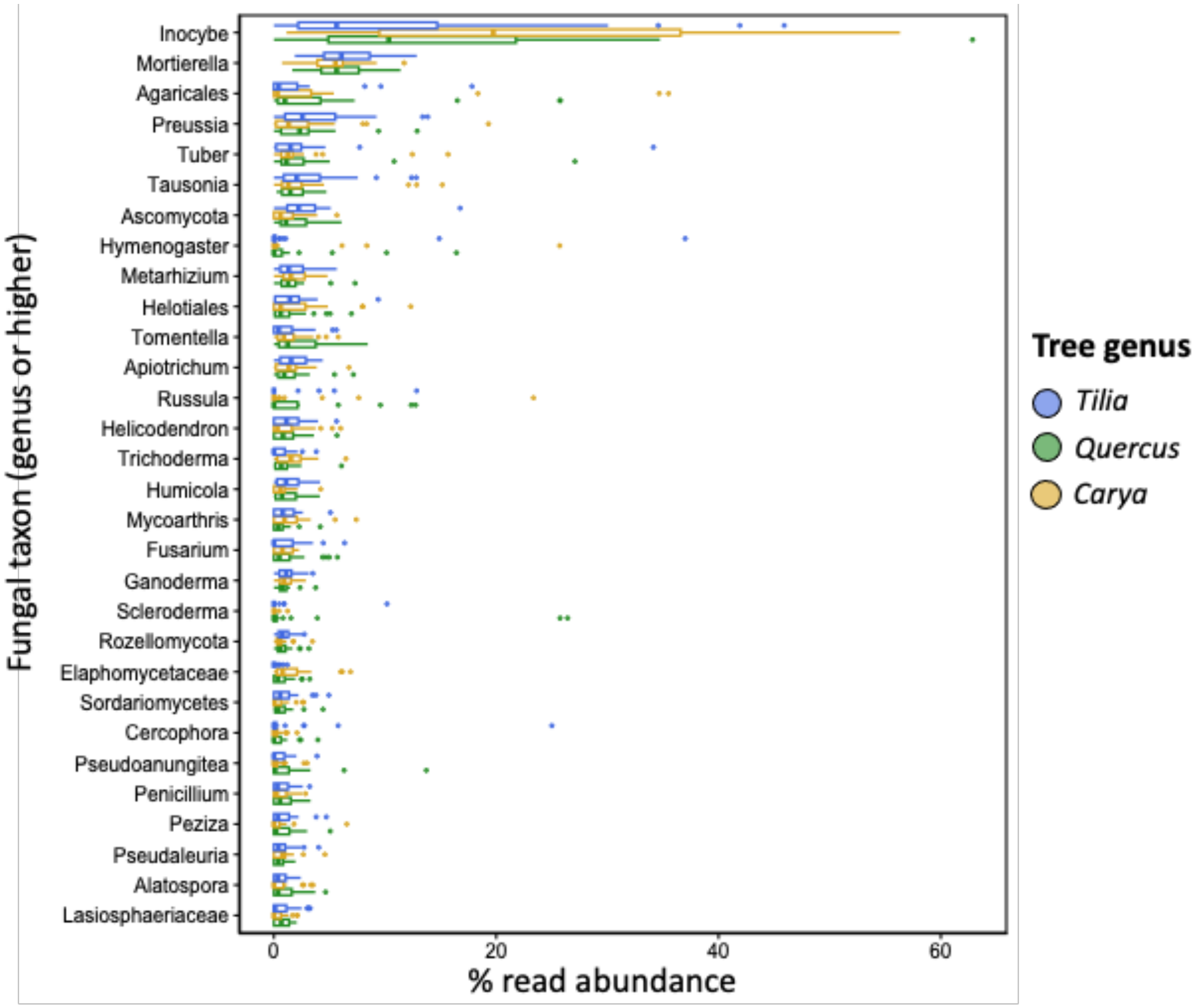


Figure S3. Relative abundance of 30 most common fungal taxa based on aggregating ASVs to lowest phylogenetic level identified at genus or higher.

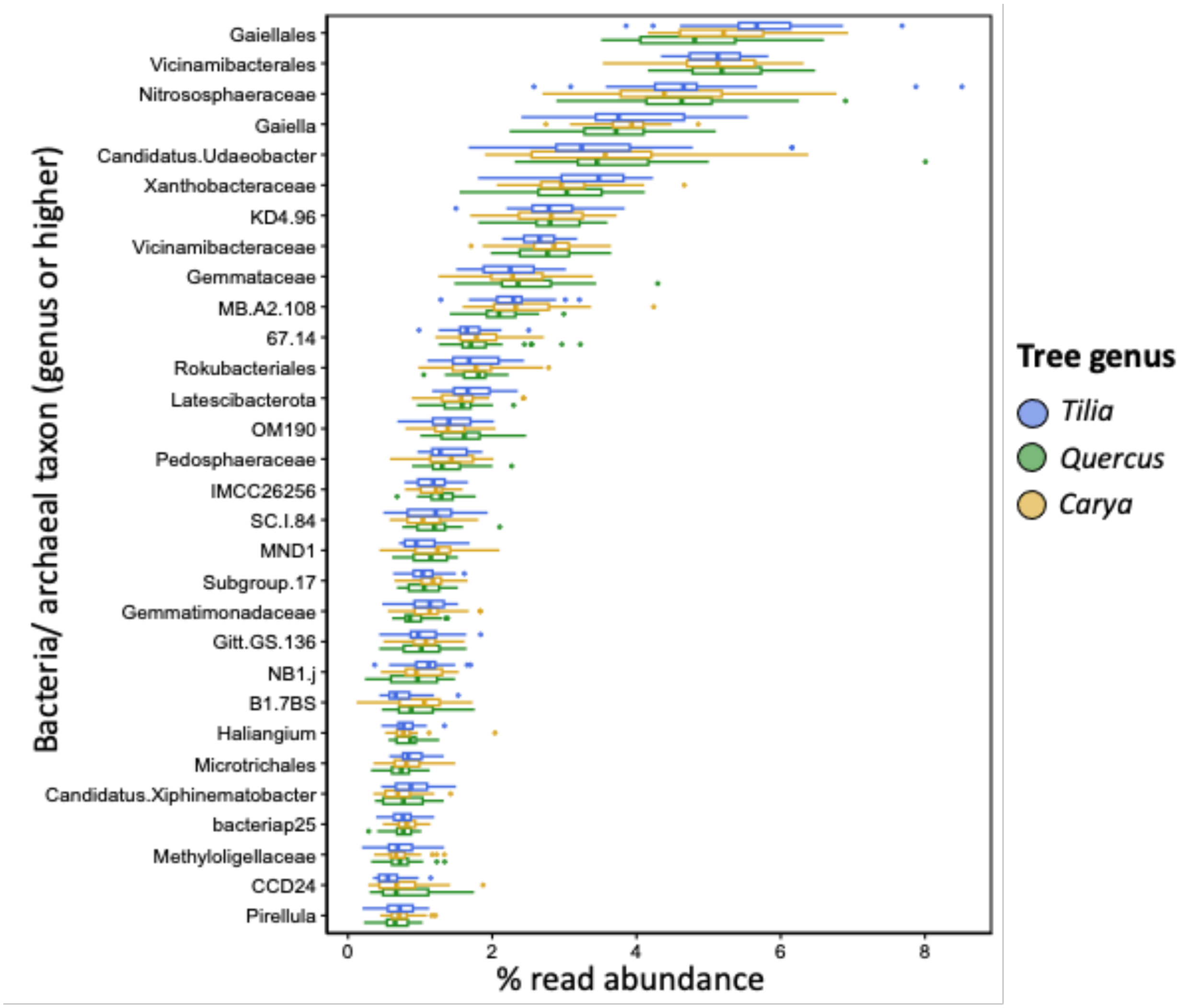


Figure S4. Relative abundance of 30 most common bacterial or archaeal taxa based on aggregating ASVs to lowest phylogenetic level identified at genus or higher.
